# Phenotypic and genotypic adaptation of *E. coli* to thermal stress is contingent on genetic background

**DOI:** 10.1101/2023.01.11.523675

**Authors:** Tiffany N. Batarseh, Sarah N. Batarseh, Alejandra Rodríguez-Verdugo, Brandon S. Gaut

**Affiliations:** Department of Ecology and Evolutionary Biology, UC Irvine, Irvine, CA, USA 92617

## Abstract

Evolution can be contingent on history, but we do not yet have a clear understanding of the processes and dynamics that govern contingency. Here we performed the second phase of a two-phase evolution experiment to investigate features of contingency. The first phase of the experiment was based on *Escherichia coli* clones that had evolved population at the stressful temperature of 42.2°C. The Phase 1 lines generally evolved through two adaptive pathways: mutations of *rpoB*, which encodes the beta subunit of RNA polymerase, or through *rho*, a transcriptional terminator. We hypothesized that epistatic interactions within the two pathways constrained their future adaptative potential, thus affecting patterns of historical contingency. Using 10 different *E. coli* Founders representing both adaptive pathways, we performed a second phase of evolution at 19.0°C to investigate how prior genetic divergence or adaptive history (*rpoB* vs. *rho*) may affect the likelihood of parallel responses and evolutionary outcomes. We found that phenotype, as measured by relative fitness, was contingent on founder genotypes and pathways. This finding extended to genotypes, because *E. coli* from different Phase 1 histories evolved by adaptive mutations in distinct sets of genes. Our results suggest that evolution depends critically on genetic history, likely due to idiosyncratic epistatic interactions within and between evolutionary modules.

## INTRODUCTION

Stephen Jay Gould famously argued that historical contingency is a defining feature of evolution (Gould 1989). He asserted that evolution is contingent on the idiosyncratic nature of historical events, leading to unpredictable and perhaps unrepeatable evolutionary outcomes. In contrast, natural selection is deterministic in the absence of historical contingency, and hence it can, in theory, converge on an optimal solution to any challenge. Although it is obvious that evolution must be historically contingent to some degree, questions remain about the patterns and mechanisms of contingency. Exploring the interplay of contingency and determinism, and the genetic effects that drive their dynamics, is crucial for understanding the evolutionary process. Moreover, understanding this interplay is helpful for predicting outcomes to applied problems - e.g., projecting which species will survive climate change, identifying chemotherapeutic agents that do not generate resistance, and forecasting pathogen variation, mutation and epidemiology (Vlachostergios & Faltas 2018; Leray et al. 2021; Bay et al. 2017).

Historical contingency has been studied in both natural populations and in the laboratory. In the field, experiments have inferred deterministic outcomes from convergent evolutionary events (Losos 2011). For example, one experiment tested brown anole lizard populations that were subjected to living on narrow perches and found that all lizard populations evolved shorter limbs (Kolbe et al. 2012). Similarly, male guppies across different populations evolved shorter life histories in the absence of predators (Reznick & Bryga 1987). These pervasive convergent outcomes have been used to argue that natural selection is predictable and that historical contingency does therefore not play a major role in evolution (Losos 2010; McGhee 2016). This interpretation is debated, however, because natural populations may contain standing genetic variation that increases the probability of parallel responses (Blount et al. 2018).

In the laboratory, controlled experiments have investigated contingency by evolving populations from a single ancestral genotype (Blount et al. 2018). For example, in the Long Term Evolution Experiment (LTEE), twelve replicated *Escherichia coli* populations converged phenotypically, as evidenced by increased fitness, faster growth, and larger cells (Bennett & Lenski 1993; Wiser et al. 2013), suggesting a lack of contingency based on mutations that arose during the experiment (Blount et al. 2018). However, one rare adaptation in the LTEE, the ability to utilize citrate (Blount et al. 2008), was contingent on previously-occurring, “potentiating” mutations. Another example from the LTEE is the evolution of antibiotic resistance, because both the level of resistance and the complement of resistance mutations varied as a function of the starting genetic background of the LTEE line (Card et al. 2021). These studies show that the evolution of phenotypes can be, but are not always, historically contingent on the genetic background. Nonetheless, important questions remain (Blount et al., 2018). For example, what circumstances favor contingent vs. deterministic evolutionary outcomes? How does prior genetic divergence between populations affect the likelihood of parallel evolutionary responses? And to what extent does epistasis shape evolutionary outcomes?

One approach to study these questions is two-phase (or ‘historical difference’; Blount et al., 2018) evolution experiments (**Figure 1**). In this experimental set-up, Phase 1 consists of initially identical populations that are evolved in the same environment, leading to potential genetic and phenotypic divergence among the populations. These populations are then subjected to a second phase (Phase 2) by allowing them to evolve in a new environment. The question is whether Phase 2 evolution differs across populations as a function of Phase 1 history. That is, is evolution in Phase 2 constrained by (or contingent upon) phenotypic and genotypic variation that was generated in Phase 1?

**Figure 1:**
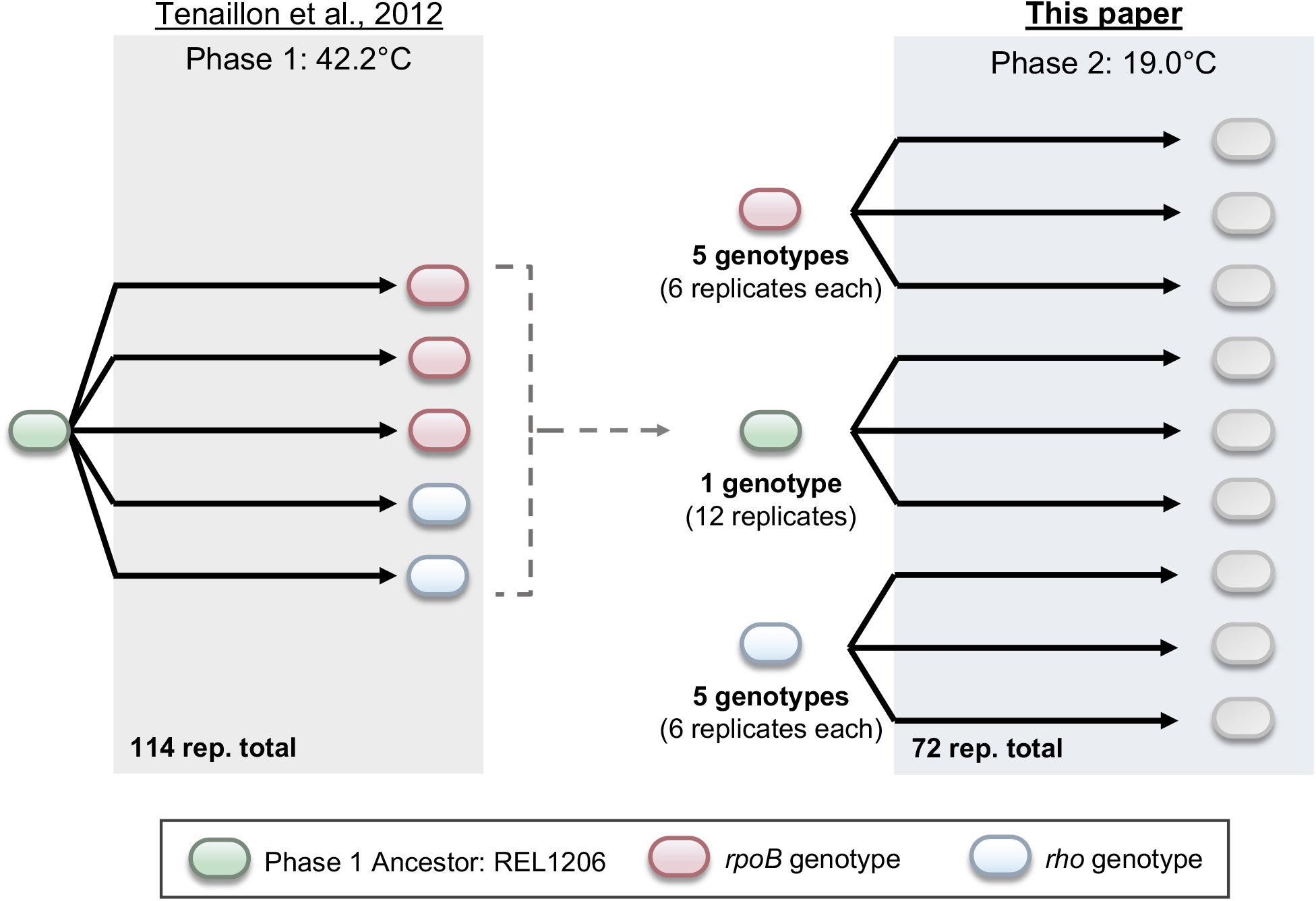
The two-phase evolution experiment designed to study contingency. The first phase of evolution was described in Tenaillon et al. (2012). The second phase used a subset of evolved close from Phase 1 and represented three pathways: clones with *rpoB* mutations (5 genotypes), clones with *rho* mutations (5 genotypes) and the Phase 1 Ancestor.

Thus far, the two-phase approach has uncovered nuanced insights into contingency. For example, Plucain et al. (2016) evolved 16 *E. coli* populations in four different chemical environments for 1,000 generations (Phase 1) before propagating them for another 1,000 generations in a single, new environment (Phase 2). The evidence for contingency was mixed: they found some evidence for contingency on phenotypic evolution, because both the growth rate and fitness of Phase 2 populations varied according to their Phase 1 history. However, they found no evidence for contingency at the molecular level; the mutations that arose during Phase 2 evolution did not vary as a function of the genetic backgrounds generated during Phase 1 (Plucain et al., 2016). Another two-phase evolution experiment used yeast and estimated that ~50% of the variance in fitness across Phase 2 populations was attributable to Phase 1 history, mostly because Phase 1 populations with low fitness evolved more rapidly in Phase 2 (Kryazhimskiy et al. 2014). However, like the *E. coli* study, this study also found no evidence to suggest that Phase 2 genotypic changes were influenced by Phase 1 history.

Based on their two-phase experiment, Kryazhimskiy et al. (2014) proposed the “global epistasis hypothesis” a form of diminishing-returns epistasis. Diminishing-returns epistasis implies that adaptive mutations have larger selective effect in relatively unfit genotypes (Griffing 1950; Jerison & Desai 2015). The global epistasis hypothesis further posits that a mutation’s effect depends solely on the fitness of the genetic background and not on the background genotype (Wei and Zhang, 2019). In this framework, evolutionary trajectories are predictable based on fitness information alone. In contrast, recent work has suggested that epistasis may be idiosyncratic, in that the direction and/or magnitude of epistatic interactions may depend on specific genotypes (Wei & Zhang 2019; Bakerlee et al. 2022), potentially making evolutionary outcomes less predictable. Idiosyncratic epistasis may even be modular, because it is a property of interactions among of genes that contribute to specific functions (Tenaillon et al., 2012) or that vary by environment (Wei and Zhang, 2019).

To examine these ideas further, we perform a two-phase evolution experiment in *E. coli* that uses extreme temperatures as the selective environments. We focus on extreme temperatures for four reasons. First, temperature is a fundamental environmental property that affects physiological traits and often defines species’ distributions; hence it often requires a complex evolutionary response (Cooper et al. 2001; Somero 1978). Second, temperature adaptation often, but not always, leads to trade-offs at other temperatures (Rodríguez-Verdugo et al. 2014), suggesting that contingency could be important in this system. Third, temperature is a topic with rich historical precedent in the experimental evolution literature. For example, Bennett and Lenski (1993) evolved *E. coli* at 20°C, near the lower edge of the temperature niche, after first adapting them to the upper end of the temperature niche (42.2°C). They found no convincing evidence of contingency at the phenotypic level, but their work was based on a relatively small number of samples (*n* = 12) and lacked genetic information.

The fourth reason that we focus on temperature is because we can take advantage of a previous large-scale experiment. Tenaillon et al. (2012) evolved 115 lines from a single *E. coli* founder strain (REL1206) at 42.2°C. After 2000 generations of evolution, they evaluated a single clone from each population for fitness gains and sequence changes. The sequence changes revealed that adaptation often occurred through two distinct adaptive pathways defined by mutations in either the RNA polymerase subunit beta (*rpoB*) gene or the transcriptional terminator (*rho*) gene. Mutations in *both* of these genes occurred statistically less often than expected by chance, suggesting negative epistatic interactions. Intriguingly, the two pathways were each positively associated with additional distinct sets of mutations. For example, *rpoB* clones tended to have mutations in *rod, ILV* and *RSS* genes, but mutations in these genes were rare in the *rho* lines. The complex landscape of both negative (e.g., *rpoB* vs. *rho*) and positive (e.g., *rpoB* with *rod* and *ILV*) epistasis suggests that evolutionary changes are partially dependent on the genetic background and that the two pathways may represent discrete evolutionary modules.

Here we hypothesize that patterns of epistasis constrain future adaptation and thus affect patterns of historical contingency. To test this hypothesis, we utilize evolved clones from Tenaillon et al. (2012) to represent Phase 1 of a two-phase experiment. After choosing a set of clones representing *rpoB* and *rho* genotypes, we evolve them at the lower extreme of *E. coli’s* temperature niche (19°C) and then measure phenotypic and genotypic differences among evolved populations. In doing so, we address three sets of questions: First, do the *rpoB* and *rho* lines differ in their response to selection at 19°C, as measured by their fitness response? We are particularly interested in testing one of the predictions of the global epistasis model, which is that populations with lower fitness should exhibit larger fitness gains. Second, is there evidence to suggest that evolution of *rpoB* and *rho* lines differ in their genotypic patterns of change? That is, do the mutations that appeared during Phase 1 evolution shape the set of adaptive mutations that accumulate in Phase 2? Finally, what do our results imply about the evolutionary process, particularly whether epistasis is global or more idiosyncratic and modular?

## RESULTS

### Selecting Phase 2 Founders that represent two adaptive pathways

Phase 1 consisted of 115 lines evolved at 42.2°C (Tenaillon et al. 2012). After 2,000 generations of evolution, single clones from these lines experienced fitness gains of ~42%, on average, relative to the single founding ancestor, which we call the Phase 1 Ancestor (**Figure 1**). To perform our Phase 2 experiment, we selected five *rpoB* clones and five *rho* clones as the founding genotypes for evolution at 19°C. We refer to these ten clones as the Phase 2 Founders (**Figure 1**), and label each by its founding line and by its *rho* or *rpoB* genotype (e.g., *rho*_A43T) (**Table 1**). It is important to recognize, however, that the Phase 2 Founders had mutations in additional genes (i.e., not just *rpoB* and *rho*) relative to the REL1206 Phase 1 Ancestor (Tenaillon et al., 2012).

**Table 1.** Phase 2 Founders.

| Phase 1<br>Evolved<br>Line* | Phase 1<br>Adaptive<br>Pathway | Phase 1<br>Adaptive<br>Pathway<br>Genotype | Mean<br>Absolute<br>Fitness at<br>19.0°C<sup>1</sup> | Mean<br>Relative<br>Fitness at<br>19.0°C<sup>2</sup> | Mean<br>Relative<br>Fitness at<br>42.2°C<sup>1</sup> | Number of<br>replicates |
| --- | --- | --- | --- | --- | --- | --- |
| 2 | *rho* | T231A | 0.018 | 0.97 | 1.484 | 6 |
| 66 | *rho* | V206A | 0.053 | 1.004 | 1.43 | 6 |
| 82 | *rho* | I15N_1 | -0.003 | 0.952 | 1.498 | 6 |
| 87 | *rho* | I15N_2 | 0.044 | 1.015 | 1.703 | 6 |
| 134 | *rho* | A43T | 0.07 | 1.008 | 1.38 | 6 |
| 3 | *rpoB* | I966S | 0.022 | 0.895 | 1.257 | 6 |
| 34 | *rpoB* | G556S | -0.082 | 0.962 | 1.51 | 6 |
| 94 | *rpoB* | E84G | 0.07 | 1.031 | 1.767 | 6 |
| 137 | *rpoB* | I966N | 0.046 | 0.982 | 1.349 | 6 |
| 142 | *rpoB* | I527L | 0.106 | 0.899 | 1.609 | 6 |
| REL1206 | Phase 1<br>Ancestor | NA | -0.004 | 1 | 1 | 12 |

\* Line numbers designated in Tenailon et al. 2012
<sup>1</sup> Relative fitness value data for 42.2°C and absolute fitness value data for 19.0°C from Rodríguez-Verdugo et al. 2014
<sup>2</sup> Relative fitness value data at 19.0°C generated for this study

We selected the Phase 2 Founders based on five criteria. First, each clone had a single mutation in either *rpoB* or *rho* but not in both genes. Second, we chose a set of clones that reflected the range of fitness values at both 19.0°C and 42.2°C for Phase 1 evolved clones (Rodriguez-Verdugo *et al*. 2014; Table 1). Third, we selected *rpoB* and *rho* lines with similar average relative fitness (*w*_r_) values at 19.0°C, at 0.954 for *rpoB* and 0.990 for *rho* (t-test, P = 0.0947). We note, however, that the *w*_r_ variance was higher among the *rpoB* founders (Var(*w*_r_) = 0.0033) compared to the *rho* founders (Var(*w*_r_) = 0.00075). Fourth, we chose a sample of distinct *rpoB* and *rho* mutations, because Phase 1 mutations occurred across different codons and caused different amino acid replacements (**Table 1**). Finally, we only chose clones that survived an initial nine-day extinction test at 19.0°C (see Methods).

Once chosen, Phase 2 Founders were propagated at 19°C for 1,000 generations, with six replicate populations per founder, under conditions identical to the Phase 1 experiment except for temperature (19°C vs 42.2°C). We also evolved 12 replicates of the Phase 1 Ancestor as a control (**Figure 1**), making a total of 72 (= 6 x 10 + 12) populations in the Phase 2 experiment. Among the 72 populations, seven went extinct, including one of the 12 descended from the Phase 1 Ancestor, four of six populations descended from Phase 2 Founder Line 3 (*rpoB* I966S), and two populations descended from Phase 2 Founder Line 142 (*rpoB* I572L) (**Table 1**). The following analyses were therefore performed on the set of 65 surviving populations.

### Relative Fitness at 19.0°C varied significantly among pathways and founder genotypes

Previous two-phase experiments have shown that fitness can be affected by historical contingency (Plucain et al., 2016; Kryazhimskiy et al. 2014). To test for such contingencies, we first measured *W_r_* of the Phase 2 populations against the Phase 1 Ancestor, based on three technical replicates per population. From a total of ~200 competition experiments, we estimated that the fitness of evolved populations increased by 3.6%, on average, at 19.0°C (P < 0.01, Wilcoxon test; **Figure 2**). On average, lines descended from *rpoB* backgrounds had 1.0% higher fitness than the Phase 1 Ancestor; the *rho* lines had higher fitness by 6.4%; and the control lines increased by 3.4% (**Table 2**). To test whether these differences were significant, we applied ANOVAs that partitioned by pathway (*rho* vs. *rpoB*) and were nested by Phase 2 founder genotypes (**Table 2**). The pathway effect was significant (p < 0.003) and explained 10.0% of the *w_r_* variance, but the Phase 2 genotype explained an even higher proportion of the variance (23.0%; p = 0.11). [Note that *w*_r_ values were not normally distributed (Shapiro-Wilk, p < 0.01), despite the large sample size and even after routine normality transformations. However, we repeated these analyses using non-parametric tests and obtained similar results (effect of pathway: p= 0.00023, η^2^= 0.0784; effect of Phase 2 Founder: p = 0.00018; η^2^= 0.153).] Overall, these observations indicate that the fitness response depended on both the pathway and the Phase 2 founding genotypes within pathways.

**Figure 2:**
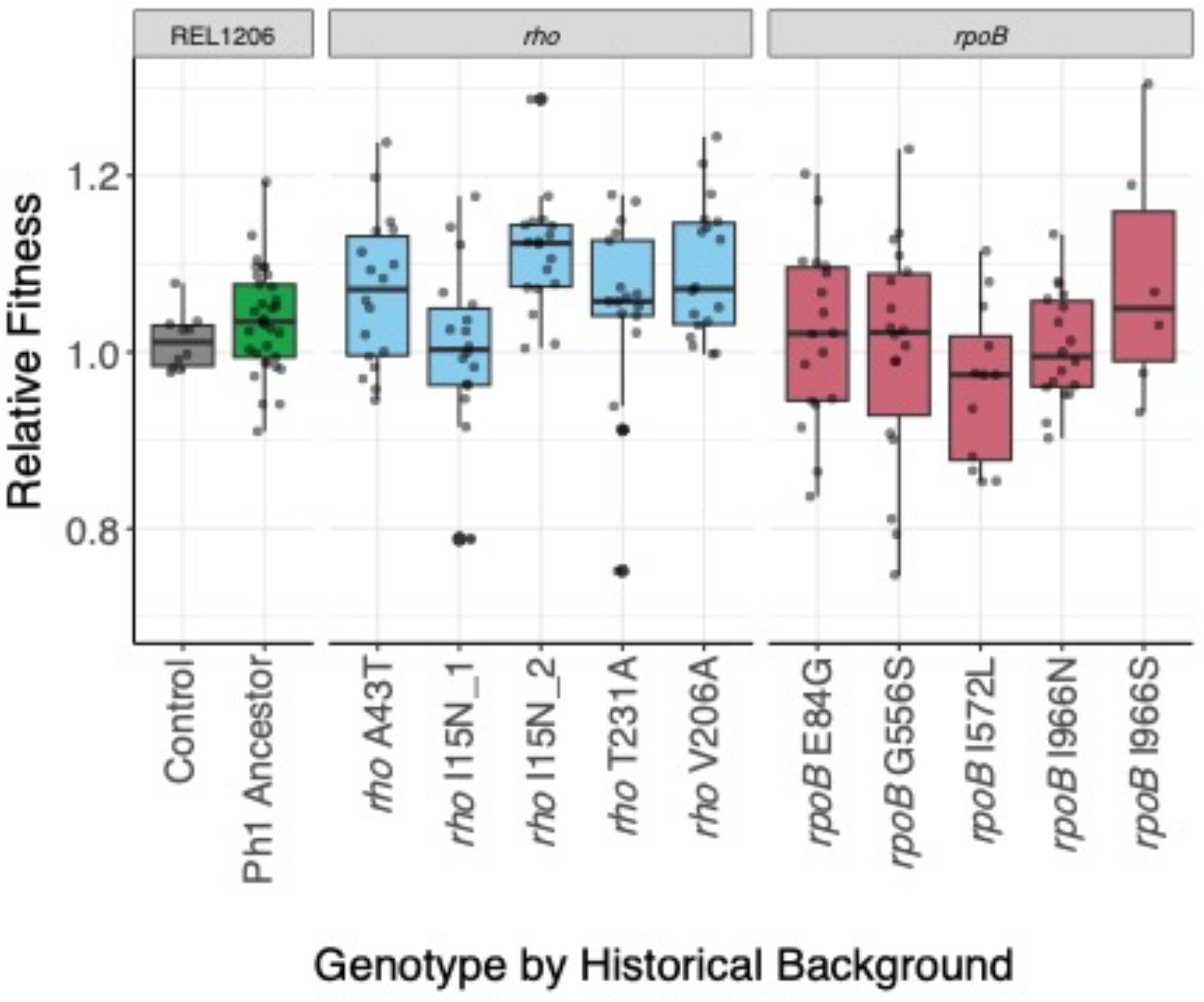
Relative fitness (*w_r_*) of the Phase 2 evolved populations at 19.0°C based on competition assays to REL1207 (an Ara+ variant of the Phase 1 Ancestor; see Methods). On the *x*-axis, Control represents the Phase 1 Ancestor competed against itself and shows that the Phase 1 Ancestor (REL1206) and the REL1207 variant have similar fitnesses, as expected. The boxplot for Ph1 Founder reports *W_r_* for the control populations evolved from REL1206, and the remaining boxplots show *W_r_* values for different Phase 2 founders. The dots in each boxplot show each *w_r_* measurement cross replicated populations, with each population measured at least three time. The horizontal line within the boxplot is the mean *W_r_* value, with the box representing the upper and lower quartile and whiskers showing the range.

**Table 2.** Relative fitness measurements for Phase 2 Evolved Populations.

| Phase 1 Evolved Line* | Phase 1 Adaptive Codon Background | Phase 1 Ancestor Competitor |  | Phase 2 Founder Competitor |  | Phase 2 Founder Competitor |  |
| --- | --- | --- | --- | --- | --- | --- | --- |
| | | Average $w_r$ (19.0°C) | P-value<sup>1</sup> (19.0°C) | Average $w_r$ (19.0°C) | P-value<sup>1</sup> (19.0°C) | Average $w_r$ (42.2°C) | P-value<sup>1</sup> (42.2°C) |
| 2 | *rho* T231A | 1.02 | **<0.01** | 1.12 | 0.17 | 0.91 | **<0.01** |
| 66 | *rho* V206A | 1.09 | **<0.01** | 1.06 | **<0.01** | 0.8 | **<0.01** |
| 82 | *rho* I15N_1 | 1 | **<0.01** | 1.04 | 1 | 0.93 | **0.02** |
| 87 | *rho* I15N_2 | 1.11 | **<0.01** | 1.13 | **<0.01** | 0.98 | 0.08 |
| 134 | *rho* A43T | 1.07 | 0.6 | 1.01 | **<0.01** | 0.96 | **<0.01** |
| 3 | *rpoB* I966S | 1.08 | **<0.01** | 1.15 | 0.2 | 1.04 | 0.44 |
| 34 | *rpoB* G556S | 1 | **<0.01** | 1.08 | 0.93 | 0.52 | **<0.01** |
| 94 | *rpoB* E84G | 1.02 | **<0.01** | 1.08 | 0.42 | 1 | 0.59 |
| 137 | *rpoB* 1966N | 1.01 | **<0.01** | 1.14 | 0.71 | 0.92 | **0.02** |
| 142 | *rpoB* I572L | 0.96 | 0.13 | 1.06 | 0.19 | 0.9 | **0.02** |
| REL1206 | Phase 1 Ancestor | 1.04 | NA | NA | **<0.01** | NA | NA |

\* Line numbers designated in Tenaillon et al. 2012
<sup>1</sup> Bolded p-values indicate a statistically significant difference from 1.0, indicating a significant change in relative fitness compared to the competitor. T-test or Wilcoxon test dependent on Shapiro-Wilk test.

In their yeast experiment, Kryazhimskiy et al. (2014) found that the rate of change of Phase 2 populations varied as a function of the fitness of Phase 2 Founders – i.e., less fit Founders led to generally larger leaps in fitness during Phase 2 evolution. To assess this potential effect, we competed Phase 2 populations against their respective Phase 2 Founders at 19.0°C, constituting another ~160 *W_r_* competition assays. On average, the Phase 2 populations had *w_r_* = 1.08, thus reflecting, on average, a significant 8% fitness advantage at the end of the experiment (P < 2.2 x 10^-16^, one-sample t-test) compared to their Founders. Lines descended from *rpoB* genotypes experienced a 9% fitness advantage on average (P = 1.41 x 10^-15^, one-sample t-test), those descended from *rho* founders had a 7% fitness advantage (P = 1.32 x 10^-10^, one-sample t-test), and the difference was not significant (P = 0.063, unpaired t-test; Figure 3A). However, the effect of the Phase 2 Founder genotype on fitness was again significant (η^2^ = 0.23, P = 1.49 × 10^-6^, Kruskal-Wallis Test), and eight of the ten sets of Phase 2 populations had significant fitness advantages relative to their Founders (**Table 2; Figure 3B**).

**Figure 3:**
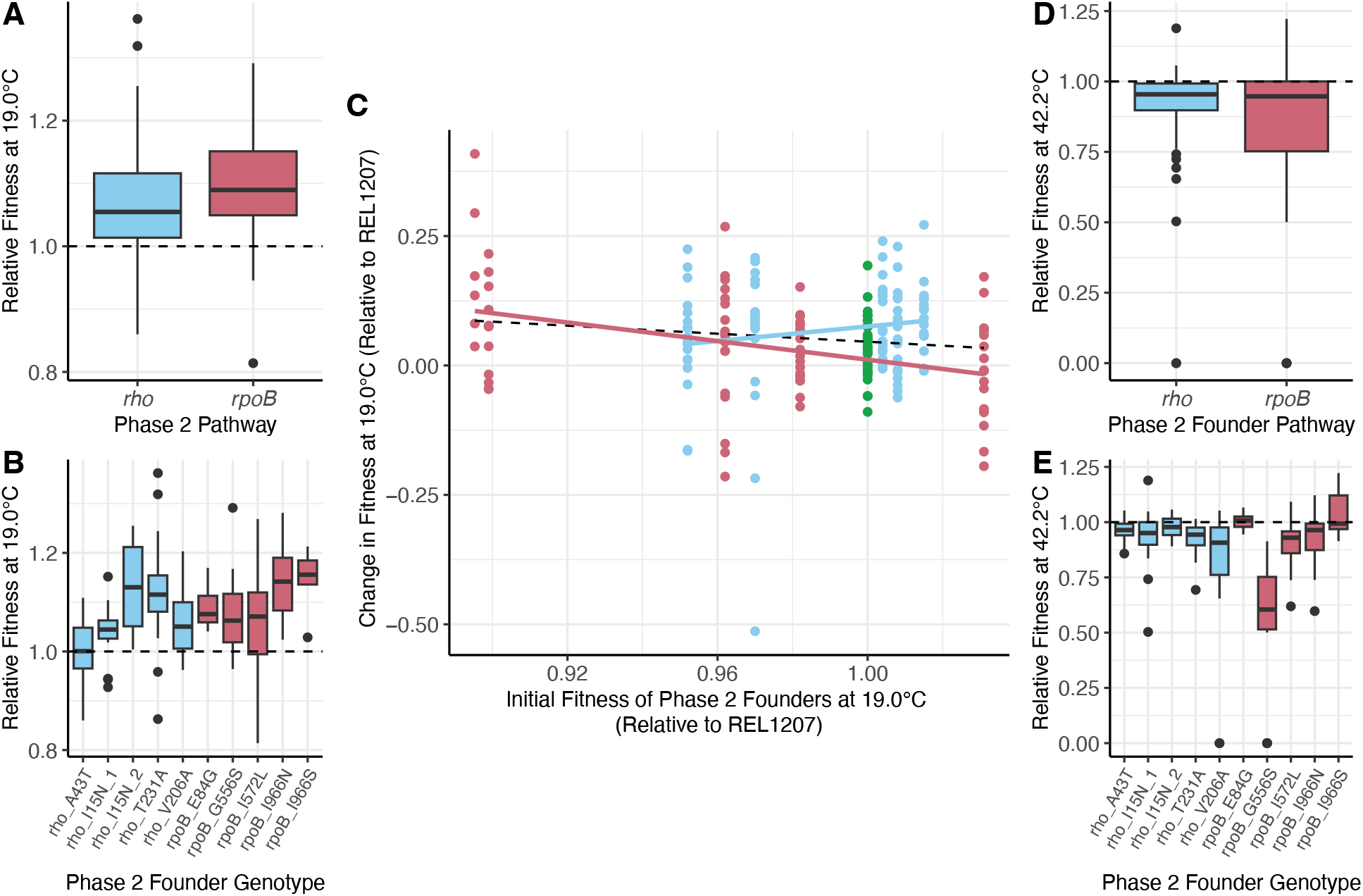
A) and B) report *W_r_* of Phase 2 evolved populations competed against their Phase 2 ancestor at 19.0°C. A) summarizes by pathways, while B) provides the information by genotype. C) The y-axis plots the change in *W*_*r* from_ the Phase 2 Founder to its evolved populations. The dots represent populations descended from the Phase 1 Ancestor (green), from *rho* genotypes (blue) and *rpoB* genotypes (red). The black, dashed line represents the overall trendline; the red and blue trendlines consider only *rpoB* and *rho* populations, respectively. D) and E) report *W_r_* of Phase 2 evolved populations competed against their Phase 2 Founder at 42.2°C, providing insights into trade-offs. D) summarizes by pathways, while E) provides the information by genotype.

To test explicitly whether populations with relatively low fitness Phase 2 Founders had larger fitness gains, we plotted *w_r_* for all Phase 2 Founders against the difference in *w_r_* between the Phase 2 Founder and each of its evolved populations (**Figure 3C**). [All of these fitness values were relative to the Phase 1 ancestor.] We expected a negative slope, and the combined set of *rpoB* and *rho* populations followed this prediction, because lower-fitness founders had slightly bigger shifts in fitness during Phase 2 evolution (slope = −0.36, Figure 3C). The trend was especially evident among the *rpoB* populations (*rpoB* slope = −0.90; Figure 3C) but did not hold for *rho* Founders and descendant populations. In fact, the slope based on *rho* populations was positive (*rho* slope = 0.70; Figure 3C). A linear model demonstrated that the two slopes were significantly different from each other (P = 0.0026). Although the cause(s) of these different patterns between pathways is not clear, it suggests, at a minimum, that the rate of fitness change during Phase 2 evolution differed by pathway and was not a simple function of the fitness of Phase 2 Founders.

### High temperature trade-offs are contingent on the founders’ adaptive history and genotype

Previous research has demonstrated significant differences in tradeoff dynamics between adaptive high-temperature genotypes (Rodríguez-Verdugo et al. 2014). For example, nearly half of the 42.2°C adapted lines from Tenaillon et al. (2012) were less fit than the Phase 1 Ancestor at lower temperatures (37°C and 20°C), slightly more than half exhibited no obvious trade-off, and a surprising few were actually fitter than the ancestor at low temperatures (Rodríguez-Verdugo et al. 2014). These trade-off dynamics imply the possibility of genotype-specific contingencies; we thus investigated trade-off dynamics at 42.2°C for Phase 2 populations relative to their Phase 2 Founders. As expected, the Phase 2 evolved populations generally had lower fitness (average *w_r_* = 0.89, P =1.14 × 10^-14^, Wilcoxon test) than their Founders at 42.2°C. Of the ten *rho* and *rpoB* Phase 2 groups, seven of ten had significantly lower *w_r_* at 42.2°C compared to their Phase 2 Founder (Table 2).

We investigated whether these patterns mapped to either pathways or starting genotypes; the difference in average *w_r_* between *rho* and *rpoB* populations was not statistically significant (P =0.38, Wilcoxon test; **Figure 3D**). It was nonetheless notable that *rho* lines experienced a fitness decline (relative to their Founder) of 8.5% while *rpoB* declined by 15% on average, suggesting some difference in trade-off dynamics between pathways. Moreover, the starting genotype had a significantly large effect on the values of *w_r_* at 42.2°C (η2 = 0.34, P = 1.36 × 10^-9^, Kruskal-Wallis test; **Figure 3E**). For example, lines descended from the *rpoB* G556S and *rho* V206A Founders had extremely low fitness at 42.2°C, at *w_r_* = 0.52 (P < 0.01, Wilcoxon test) and *w_r_* = 0.80 (P < 0.01, Wilcoxon test). The take-home point is that trade-offs differed as a consequence of Phase 2 Founder pathways and genotypes.

### Mutations that arose during Phase 2 evolution are contingent on the genetic history

To examine contingency at the level of individual mutations, we sequenced the DNA of all 65 Phase 2 and control populations. After filtering the sequencing data and calling genomic variants, we identified 1,387 point mutations and short indels (<50 bases in length) that arose during the Phase 2 experiment and had population frequencies > 5% (Supplemental Figure 1). Almost half (45%) of the 1,387 mutations were present at a frequency of < 10%, but 119 were “fixed” (as defined by a frequency > 85% in single evolved population). Fixed mutations were present in 51 of the 65 sequenced populations, but there was no discernible pattern across the 14 lines that lacked fixed mutations by pathway (*rho*, *rpoB* and control Founders). Overall, the largest proportion of mutations 54.4% (742/1387) occurred in intergenic regions (Figure 4A), and 95.8% of these were point mutations. Within genes, most (89.8% or 371/413) point mutations were nonsynonymous (371/413).

**Figure 4:**
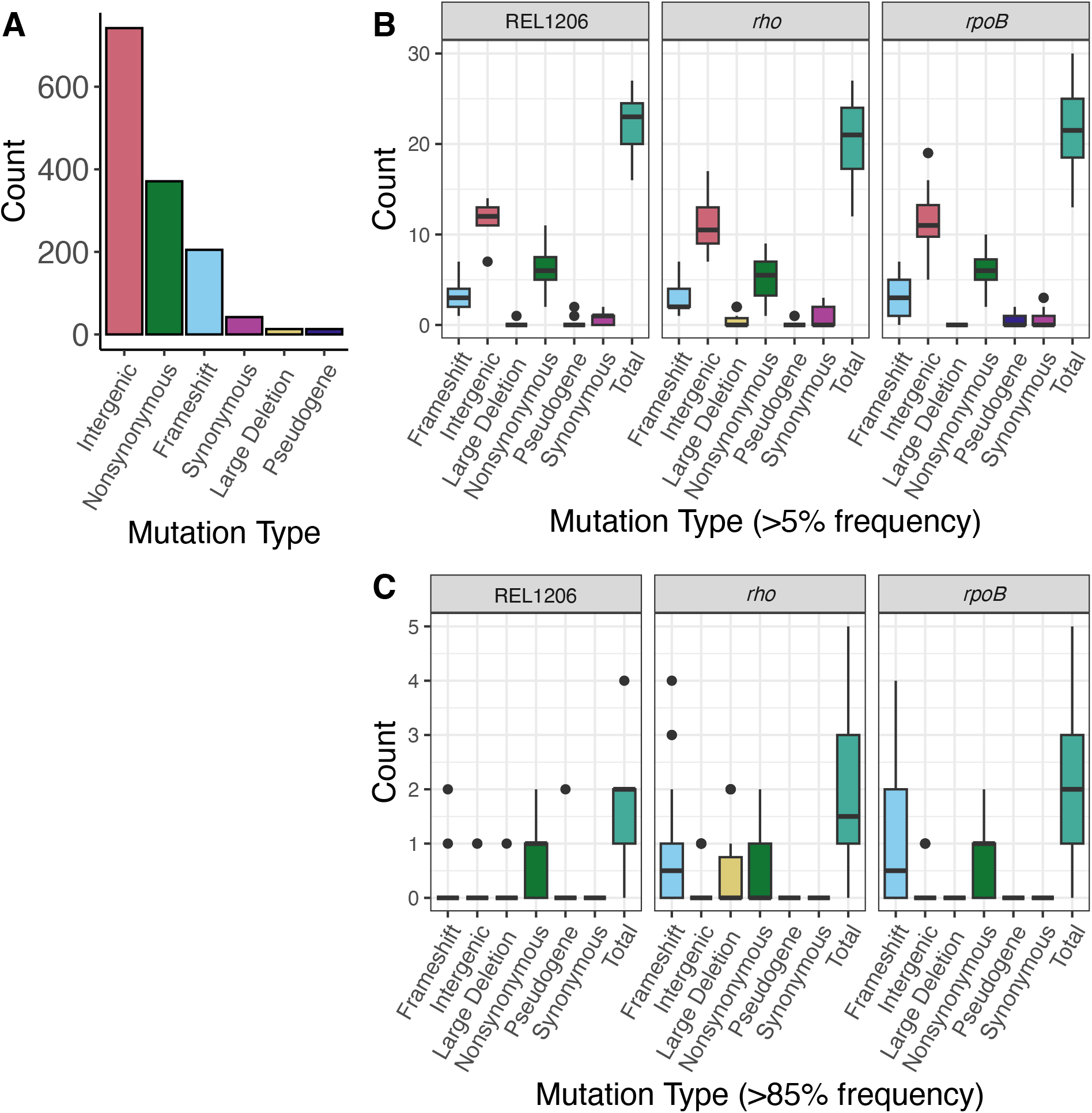
Mutations that arose during Phase 2 evolution across populations. A) Number of mutations by type across all evolved populations. B-C) Types of mutations in Phase 2 Evolved Populations faceted by pathway (Phase 1 Ancestor/REL1206, *rho* or *rpoB*). B) All mutations at 5% frequency or higher and C) fixed mutations at 85% frequency. The x-axis shows mutation types, including mutations in pseudogenes and large deletions (deletions greater than 50 bp).

We used the sequencing data to confirm that our populations were not cross contaminated during the 1,000 generation experiment by first assessing whether mutations from the Phase 2 Founder were fixed in the evolved populations, as expected. It was true in every case. We then built phylogenies based on all of the sites in the Phase 2 populations that differed from the Phase 1 Ancestor. The data confirmed expected phylogenetic relationships based on the experimental design (Supplemental Figure 2) and thus yielded no evidence of contamination.

Given a lack of obvious evidence for contamination, we first asked whether the patterns and numbers of mutations differed significantly among pathways. We identified 22 Phase 2 mutations in *rpoB* lines and 20 Phase 2 mutations in *rho* lines that were at a frequency of 5% or higher in the population on average, a difference that was not significantly different (P = 0.17, unpaired t-test). There was also no difference between pathways in the number of fixed mutations (P = 0.14, unpaired t-test). We also contrasted the proportions of mutational variant types (intergenic, frameshift, nonsynonymous, and synonymous mutations and large deletions > 50 bps) between *rho* and *rpoB* pathways, again finding no difference for the complete set of mutations (P = 0.78, contingency test; Figure 4B) or for fixed mutations (P = 0.0803, contingency test; Figure 4C). Thus, we detected no obvious difference in the number or pattern of mutations between *rho* and *rpoB* mutations.

We then investigated whether there was evidence of historical contingency at the level of specific mutations. Did *rho* and *rpoB* lines tend to accumulate mutations in different sets of genes? We first used a phylogenetic approach: focusing only on mutations that arose during Phase 2 evolution, we calculated a distance matrix and resulting Neighbor-Joining tree from the presence-absence of mutations among populations. We then tested for associations between phylogenetic clustering and the pathways of origin (i.e., *rho*, *rpoB* or control lines evolved from the Phase 1 Ancestor) against the null hypothesis of no associations between pathway and phylogenetic history. We found significant association between the mutations that arose during Phase 2 and the adaptive history at the level of pathway (ANOSIM R = 0.139, P = 4 × 10^-4^; Figure 5A). We also we tested the association between mutational patterns and variation across the Phase 2 Founder genotypes, instead of pathways. Here again, the test was significant (ANOSIM R = 0.2398, P = 1 × 10^-4^). These results are consistent with the idea that the identity of mutations differed among Phase 2 populations based in part on their Founder genotype.

Our second approach relied on Dice’s similarity coefficient (DSC) (Dice 1945). Following Card et al. (2021), we calculated DSC at the genic level between all pairs of evolved populations. We based DSC on two sets of mutations: all Phase 2 mutations (Supplemental Figure 3) and Phase 2 fixed mutations (Figure 5B). For the former, the average DSC between populations was 0.38, indicating that the evolved populations shared 38% of their mutated genes on average. The mean DSC within a historical background was 0.39, which was similar to, but significantly different from, the mean DSC between historical backgrounds (DSC=0.37; P = 4.87 x 10^-14^, Wilcoxon Test). The same analyses based on fixed mutations had an average DSC 0.10 across all comparisons, an average DSC of 0.18 between populations from different pathways, and a mean DSC of 0.06 within the same pathway (**Figure 5C**). The average DSC within and between historical backgrounds again differed significantly (P < 2.2 x 10^-16^, Wilcoxon Test), suggesting that fixed (and presumably adaptive) mutations differed among populations in part due to their Phase 2 Founder. We also detected a significant, moderate effect of the adaptive history on DSC across comparison types (η2 = 0.078, P < 2.2 x 10-16, Kruskal-Wallis test). For example, *rpoB* evolved populations had higher DSC values to each other than to populations derived from *rho* or the Phase 2 Founders (P < 0.5, Post-Hoc Pairwise Wilcoxon/Mann-Whitney test). These results further indicate that the fixed mutations that arose in the Phase 2 experiment differed due, in part, to genetic history.

**Figure 5:**
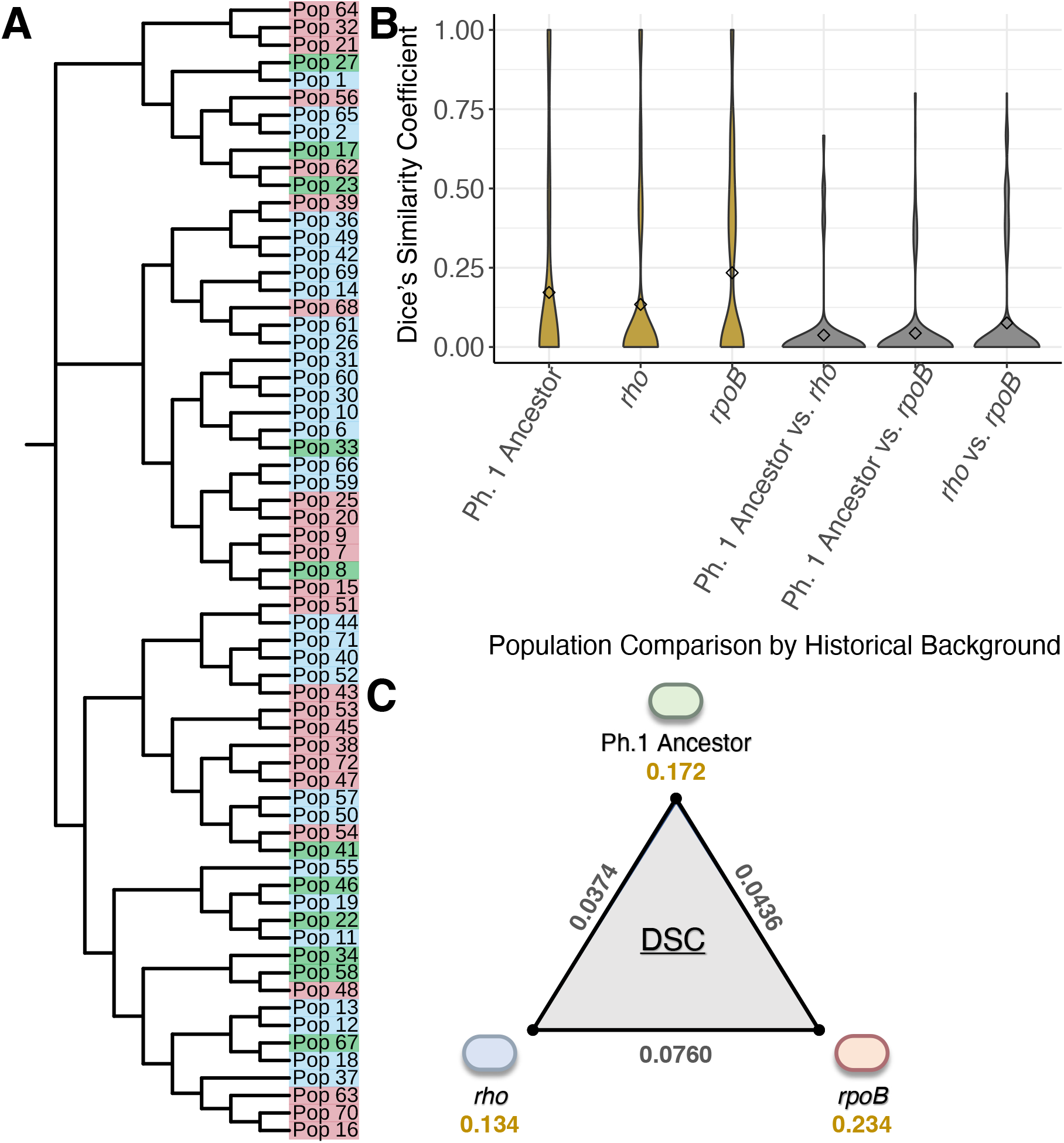
Measures of association between the evolved populations, their historical backgrounds, and the mutations that arose during Phase 2 evolution. A) A neighbor-joining tree built from presence-absence patterns of mutations that arose in Phase 2 evolved populations. Populations descended from the Phase 1 Ancestor are colored in green, with *rho* derived populations in blue, and *rpoB* derived populations in red. B) Dice’s similarity coefficients calculated from fixed mutation data, separated by the type of pairwise comparison. Pairwise comparisons performed within a pathway are in gold and those between pathways in grey. C) Average DSC values within and between historical pathways. The average between-pathway values DSC is plotted along the edges in grey, and the average within-DSC are depicted in gold.

Finally, we sought to identify specific genes that differed in their propensity to house mutations in *rho* vs. *rpoB* pathways. To do so, we counted the number of populations with and without a mutation in each gene or intergenic region and performed these counts separately for *rho* and *rpoB* populations. Using a Fisher’s Exact Test (FET), we identified six genes or intergenic regions that were more frequently mutated in one adaptive pathway but not the other (P < 0.05, Table 3). The significance levels of these tests were no longer significant (P > 0.05) after FDR correction, likely reflecting low statistical power due to sample size and many (163) FET tests. Nonetheless, the results provide evidence of contingency that affects specific genomic regions.

**Table 3.**
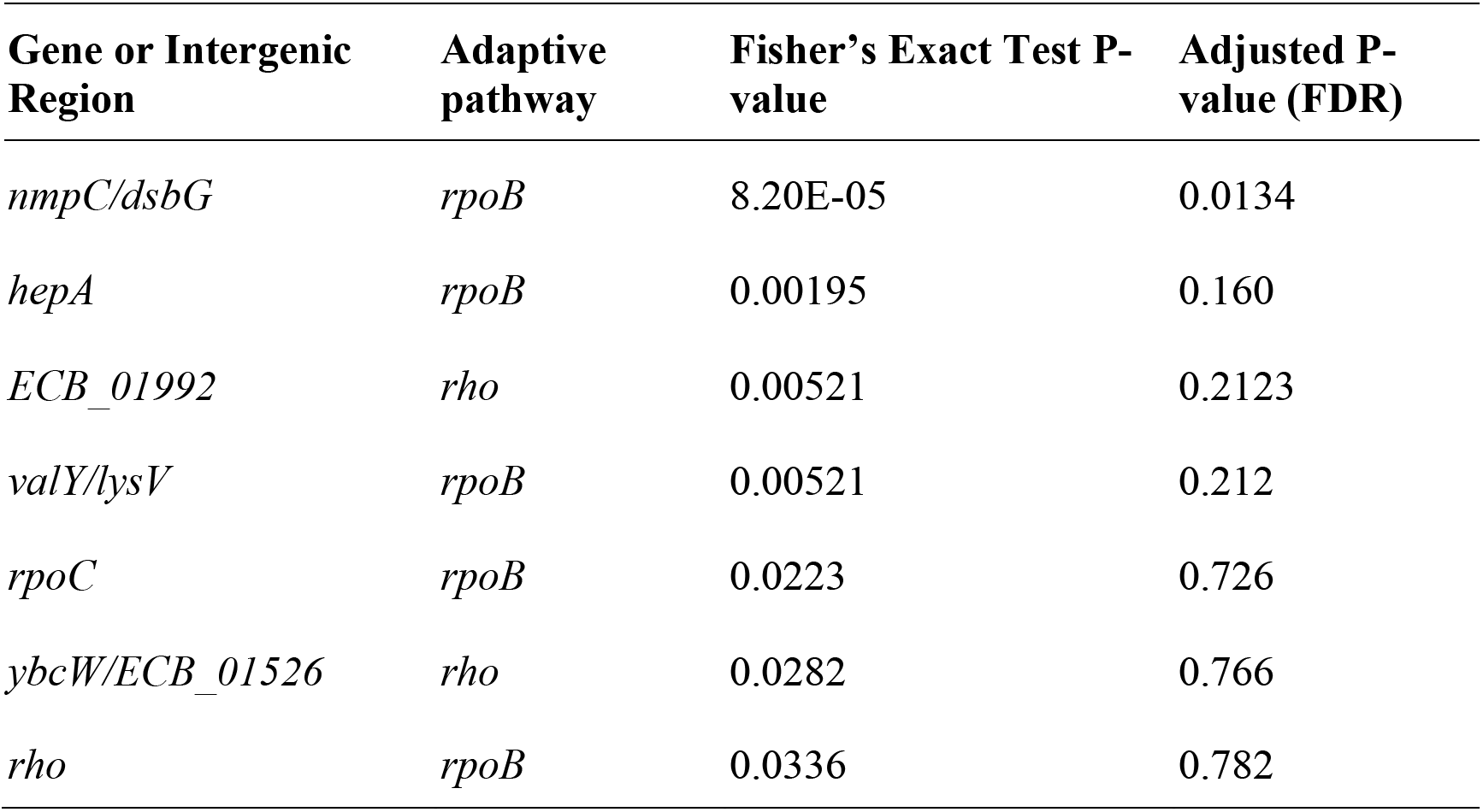
Genic or intergenic regions with evidence of biased mutation histories by pathway.

## DISCUSSION

Evolution is an inherently historical process, but the magnitude and effect of history on adaptation remains somewhat enigmatic. To yield insights into the dynamics of historical contingency, we have performed the second phase of a two-phase evolution experiment. Phase 1 was based on a study that evolved 115 initially identical populations of *E. coli* to the stressful temperature of 42.2°C (Tenaillon et al. 2012). These populations evolved primarily by one of two distinct pathways involving mutations in either the RNA polymerase beta subunit gene (*rpoB*) or the transcription termination factor *rho*. We chose five clones from each of the two pathways and evolved them for 1,000 generations in a second, low temperature (19.0°C) environment. At the end of the Phase 2 experiment, we compared the phenotypes (relative fitness, *Wr*) and genotypes of evolved populations to both their immediate ancestors (Phase 2 Founders) and to the ancestor of the entire experiment (Phase 1 Ancestor; **Figure 1**).

Based on these data, our first finding is that *W_r_* varied significantly among evolved populations both among pathways (*rho* or *rpoB*) and among Founder genotypes (**Figure 2**), explaining 10% and 23% of the *W_r_* variance. Thus, the fitness of Phase 2 populations was contingent upon their Founder fitnesses and genotypes. A more nuanced question is whether there was a predictable pattern to the fitness response. It is reasonable to expect, based on diminishing-returns epistasis, that *W_r_* among populations vary as a function of the fitness of the Phase 2 Founder, specifically that lower-fitness Founders give rise to populations with more dramatic fitness gains. We have found the expected general trend across all *rho* and *rpoB* Phase 2 populations (**Figure 3C**) and for the *rpoB* lines. Surprisingly, however, this relationship did not hold for the *rho* lines (**Figure 3C**). These contrasting results not only suggest differences among pathways (**Figures 3AB**) but raise important questions about what might drive these differences.

One must first consider the caveats and limitations of our experimental design. For example, practical considerations limited the number of *rho* and *rpoB* Founders; perhaps more or different *rho* samples would have yielded additional insights. Moreover, one of the *rpoB* Founders (*rpoB* I966S) had a much lower *w_r_* than the rest of the Phase 2 Founders, experienced the biggest shift in fitness during Phase 2 fitness, and may have driven the overall trend (**Figure 3C**). Unfortunately, we did not have a Phase 2 *rho* founder for comparison, because the only potential Phase 2 *rho* founders with similarly low fitness did not survive a 9-day extinction test. Finally, we recognize that the timescale of 1,000 generations likely does not provide a complete view of the fitness landscape; perhaps populations will converge on the same fitness optima with further evolution.

Another explanation has to do with the structure of epistasis. The ongoing debate about diminishing-returns epistasis does not center on whether it exists, because diminishing-returns have been found both by testing the effects of individual mutations (Moore et al. 2000; Kryazhimskiy et al. 2009; Perfeito et al. 2014) and by inferring patterns of evolutionary change for experimental evolution data (Khan et al. 2011; Wang et al. 2018; Bakerlee et al. 2022; Chou et al. 2011). Instead, it centers on whether the dynamics of diminishing-returns can be predicted based on starting fitness alone or whether diminishing-returns is a product of more idiosyncratic epistatic interactions that depend, in part, on the genetic background. As an example of the latter, Card et al. (2019) measured the evolution of antibiotic resistance and found that less-fit genotypes do not always evolve bigger changes in fitness resistance. Like Card et al. (2019), our experiment has not been designed to measure diminishing-returns epistasis directly. Nonetheless, our results suggest that fitness evolution is more idiosyncratic than predicted by the global epistasis model, given differences in fitness responses between pathways and genotypes (**Figures 2, 3ABC**). Several recent studies have similarly concluded that epistasis is often idiosyncratic (Wei & Zhang 2019; Bakerlee et al. 2022; Lyons et al. 2020), calling to question the universality of the global epistasis model.

The next pertinent question is: What might drive these idiosyncrasies? We do not have a complete answer to this question, but we can offer some insights. Previous work has shown that the complete set of 115 high-temperature adapted lines differed substantially in their fitness trade-offs between 42.2°C and 19°C (Rodriguez-Verdugo et al., 2014, 2017). A few lines that evolved at 42.2°C were *more* fit than their ancestor at 18°C, while others were much less fit. Our work further illustrates that Phase 2 populations vary in their trade-off dynamics (**Figure 3DE**). Furthermore, studies of single mutants have shown that some of the *rpoB* mutations in this study confer fitness advantages at 42.2°C, but in contrast two *rho* mutations (*rho* A43T and *rho* T231A), likely require positively epistatic interactions to become adaptive (González-González et al. 2017). We suspect that all of these patterns feed into idiosyncratic evolutionary responses. Given, for example, that some genotypes appear to be thermal specialists and others are generalists, one can envision that founding populations with distinct generalist *versus* specialist mutations have substantially different numbers, directions, and types of potential epistatic interactions across the genome.

A unique feature of our work is that the set of Phase 2 Founders represent pathways that were defined by epistatic interactions among distinct set of mutations and genes. We predicted that these pathways affect the evolutionary response by shaping the type and identity of future mutations. Although the two pathways do not vary in their number or types of mutations (**Figure 4**), there is ample evidence to support our prediction. For example, Phase 2 mutations cluster non-randomly on a phylogeny (**Figure 5A**), suggesting that the set of successful mutations is not independent of the Phase 2 Founding genotype. Similarly, the complement of Phase 2 mutations is more similar within a pathway than between pathways (**Figure 3B**). Finally, specific genic and intergenic regions vary in their enrichment for mutations depending on the genetic pathway of their Founders (**Table 3**). [We included intergenic regions because they have been previously implicated as drivers for bacterial adaptation (Khademi et al. 2019).] These patterns hold to some extent for the entire complement of > 1,000 mutations, but they are especially clear for the set of 119 fixed mutations (**Figure 5**; **Table 3**). Since fixed mutations are more likely to be adaptive, and also because the number of mutations is unlikely to be limiting in this system, our results show that the identity of adaptive mutations depends on genetic background, likely due to idiosyncratic interactions.

Although our results are not compatible with the global epistasis model (Kryazhimskiy et al. 2014), they do appear to adhere to a modular model of evolution (Tenaillon et al., 2012; Wei and Zhang, 2019). Of course, two-phase evolution experiments do have inherent biases, as the second phase is always founded by lines already defined by differences in evolutionary outcomes. In this case we have introduced an additional bias, because we chose Founders from two distinct pathways. However, this should not inhibit our ability to distinguish between the global or modular models of epistasis in the second phase of evolution. Interestingly, Wei and Zhang (2019) have documented that an emergent property of their modular model is the appearance of global diminishing-returns epistasis (Wei and Zhang, 2019). That is, epistatic interactions within specific modules may combine, in some cases, to provide an apparent signal of global diminishing-returns epistasis. This emergent property may explain why diminishing-returns epistasis sometimes appears to be a genome-wide phenomenon but perhaps more often does not.

Our finding -- i.e., that mutations detected in Phase 2 mutations are associated with Founder genotype -- may provide insights about mechanisms of adaptation. We have identified seven genes and intergenic regions that are enriched for Phase 2 mutations in either *rpoB* or *rho* populations (Table 3). Of the seven, five are more likely to accrue mutations within *rpoB* populations. In this context, it is worth recalling that *rpoB* is component of RNA polymerase, which is a global regulator of gene expression and that has numerous pleiotropic effects and potential epistatic interactions. At least three of the five enriched regions are related to transcriptional function: *rpoC* also codes for beta subunit of RNA polymerase (Conrad et al. 2010; Trinh et al. 2006); the RHO protein terminates RNA polymerase activity; and *hepA (which* also known as *rapA*) codes for a transcription factor with ATPase activity and also an RNA polymerase associated protein (Sukhodolets et al. 2001). Previous work has shown that modifying RNA polymerase is a key feature of adaptation to thermal stress but also that this is a blunt instrument that may cause more phenotypic changes (as measured by gene expression; Rodriguez-Verdugo et al., 2017) than may be necessary to achieve fitness gains. If true, it is reasonable to speculate that adaptation to 19.0°C from 42.2°C includes further tuning of RNA polymerase, as reflected by an enrichment of genes related to transcriptional activity like *rpoC, rho* and *hepA*.

Three further features about the enriched regions stand out. First, two enriched regions within *rpoB* populations (*valY* and *lysV;* Table 3) encode tRNA synthetases (Andersen et al. 1997; Ruan et al. 2011; Agrawal et al. 2014), suggesting that one additional or alternative route to adaptation is through modifications of translational speed or dynamics. Second, the Phase 2 evolved populations that descended from *rho* backgrounds were significantly enriched in one gene (*ECB_01992*) and one intergenic region (*ybcW/ECB_01526*) (Table 3). Both of these regions have unknown functions, and thus they yield no clues into the molecular mechanisms of 19.0°C adaptation for *rho* populations. Finally, it is interesting to speculate about the fact that more enriched regions (5 vs. 2) were found in *rpoB* vs. *rho* lines. Previous work has shown that engineered mutations of *rho*_A43T and *rho*_T231A lead to fewer modifications of gene expression than do *rpoB* mutations I572L, I572N and I966S (Gonzalez-Gonzalez et al., 2017). These observations suggest that some of the *rho* mutations are “less-connected” than the *rpoB* mutations, which could again lead to substantially different dynamics of fitness and epistsis between the two pathways.

To sum, our Phase 2 experiment has shown that the fitness response of evolved population varies by the pathway and phenotype of their Founders, but Founder fitness is not strongly predictive of fitness gains. More importantly, our work clearly demonstrates that the suite of fixed and presumably adaptive mutations in Phase 2 differs according to their Phase 1 history. Our observations indicate that history has shaped, defined, and perhaps even canalized the adaptive response of Phase 2 populations. Overall, these observations are not consistent with the global epistasis hypothesis, but instead add to a growing literature suggesting both that epistasis is idiosyncratic and evolution is often contingent on genetic history.

## METHODS

### Two-Phase Evolution Experiment Isolate Criteria and Selection

To study evolutionary contingency, we chose ten clones from Tenaillon et al. (2012) (**Table 1**) to evolve at 19°C, which is towards the lower limit of the temperature niche for the REL1206 ancestor (Rodríguez-Verdugo et al. 2014). It is worth noting that the REL1206 ancestor had been evolved for 2,000 generations at 37.0°C in minimal media prior to the Phase 1 experiment and was therefore preadapted to media and laboratory conditions (Lenski et al. 1991).

Before clones were subjected to the Phase 2 evolution experiment, they were first assessed for survivability. To test survivability, isolates from frozen stock were placed into Luria-Bertani medium (LB) and incubated at 37.0°C for one day to acclimate from frozen conditions (Rodríguez-Verdugo et al. 2014; Bennett & Lenski 1993; Lenski & Travisano 1994). The overnight culture was diluted 1,000-fold in saline and this dilution was transferred into fresh Davis Minimal (DM) Media supplemented with 25mg/L of glucose and grown for one day at 37.0°C. Following incubation, 100 μl of the culture was transferred into 9.9ml of fresh DM media and incubated at 19.0°C and serially propagated for at least nine days. Each day we measured the cell density to determine if extinctions had occurred, by diluting 50 μl of overnight culture into 9.9ml of Isoton II Diluent (Beckman Coulter) and measuring cell density in volumetric mode on a Multisizer 3 Coulter Counter (Beckman Coulter). An isolate survived if its cell density measurements were maintained over the course of the test while allowing for fluctuations of +/− 1×10^6^ cells.

### Evolution Experiment at 19.0°C

To prepare the isolates for the Phase 2 experiment, the Phase 2 Founders and the Phase 1 Ancestor (REL1206) were grown from frozen stock in 10 ml of LB at 37.0°C with 120 RPM. After 24 hours of incubation, the overnight cultures were diluted 10,000-fold and plated onto TA plates and incubated at 37.0°C. On the next day, single colonies were picked from the plates and inoculated into 10 mL of fresh LB and incubated at 37.0°C with 120 RPM. The next day, we transferred 100 μl of the bacterial culture into 9.9 ml of fresh DM25 media, which was incubated at 37.0°C at 120 RPM for 24 hours to acclimate to experimental conditions, following common practice (Lenski & Travisano 1994; Bennett & Lenski 1993; Rodríguez-Verdugo et al. 2014). After incubation, we began the Phase 2 evolution experiment by transferring 100 μl of culture into 9.9 ml of fresh DM25 and incubated the tubes at 19.0°C with 120 RPM and incubated for 24 hours.

Each day, the cultures were transferred daily into fresh media via a 100-fold dilution. At regular intervals (at generation 100 and roughly every 200 generations after that), we mixed 800 μl of each line with 800 μl of 80% glycerol to prepare whole population frozen stocks. We began the experiment in January 2020 but had to pause it after 297 bacterial generations due to the Covid-19 pandemic. To restart the experiment, we revived the bacterial populations by transferring 100 μl of thawed glycerol stock into 9.9 mL of fresh DM25 media and continued the experiment until the bacteria had grown for 1000 generations or 152 days.

### Measuring Relative Fitness

We performed competition experiments to measure the relative fitness of the Phase 2 evolved lines. We competed the Phase 2 evolved lines against the Phase 1 Ancestor at 42.2°C and their respective Phase 2 Founders at both 19.0°C and 42.2°C. To perform the competitions, we mixed the cells in a single glass culture tube and plated the mixture to count the colonies before and after 24 hours of competition. We used the neutral Ara+ marker to differentiate between the two lines when plating on tetrazolium-arabinose (TA) plates. To generate Ara+ mutants from the Phase 2 Founders for competitions, we followed previously published methods (Lenski et al. 1991). To validate neutrality, we competed the Ara+ mutants against the original Ara-stock using the methods described below. Control competition experiments were performed by competing the Phase 1 Ancestor, *E. coli* strain B REL1206, against its Ara+ mutant, REL1207 (Wiser & Lenski 2015).

To perform competition assays, bacteria from frozen glycerol stocks were revived with a loop into 10 mL of LB and incubated at 37°C with 120 RPM for 24 hours. After incubation, the overnight cultures were vortexed and 100 μl of each were diluted in 9.9 ml of 0.0875% saline solution. From each dilution tube, 100 μl was transferred to 9.9ml DM25 to incubate at 37.0°C with 120 RPM for 24 hours. Following incubation and in order for the bacteria to acclimate to the experimental temperature, we transferred 100 μl of the overnight cultures into 9.9 ml of DM25 and incubated the tubes at the experimental temperature (19.0°C or 42.2°C) with 120 RPM for 24 hours (Bennett & Lenski 1993). The next day, we mixed the Ara- and Ara+ competitor strains into sterile DM25 media. For competitions at 19.0°C, we mixed the bacteria 1:1. For competitions at 42.2°C, we mixed the bacteria 1:1 or we adjusted the ratio to 1:3 if the original ratio resulted in too few colonies (<20) on the plate for either competitor. The mixture was incubated at the experimental temperature with 120 rpm for 24 hours. After allowing the cells to compete, we quantified the cell density of each competitor by plating the overnight culture onto tetrazolium-arabinose (TA) plates and counting the number of colonies. All competitions were performed in at least triplicate, resulting in roughly 600 competitions.

Using the methods described in Lenski *et al*. (1991) and Tenaillon *et al*. (2012), we calculated the relative fitness, *W_r_*. The fitness of a Phase 2 evolved line relative to its competitor was estimated by:

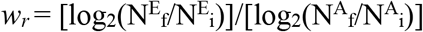

where E refers to the evolved line and A refers to the ancestral clone, where N^E^_i_ and N^A^_i_ represent the initial cell densities of the two competitors, and N^E^_f_ and N^A^_f_ represent the final cell densities after one day of competition.

### DNA Library Preparation and DNA Sequencing

To sequence the evolved populations, we revived populations from ~10 μl of frozen glycerol stock in 10ml of DM media supplemented with 1000mg/L of glucose. The culture tubes were incubated at 19.0°C with 120 RPM. We extracted from 65 bacterial populations using the Promega Wizard Genomic DNA Purification kit. DNA concentrations were measured with Qubit dsDNA HS Assay kits. We prepared our DNA sequencing libraries with the Illumina Nextera DNA Flex Library Preparation kit. The libraries were multiplexed and sequenced using the Illumina NovaSeq on an S4 flow cell to generate 100bp paired-end reads at UC Irvine’s Genomics High-Throughput Facility (https://ghtf.biochem.uci.edu). Sequencing read quality was assessed with FastQC v. 0.11.9 (http://www.bioinformatics.babraham.ac.uk/projects/fastqc), trimmed with fastp v. 0.23.2 (Chen et al. 2018), and visualized with MultiQC v. 1.9 (Ewels et al. 2016). Each population had 25,000,000 sequencing reads on average (min = 8,600,000 reads, max = 35,000,000 reads), resulting in a minimum of > 150x coverage per population.

### Variant Detection

We detected mutations and their respective frequencies in each evolved Phase 2 population using breseq v. 0.35.5 (Deatherage & Barrick 2014). We performed the breseq analysis in polymorphism mode with two different reference genomes. First, we performed breseq analysis using *E. coli* strain B REL606 as the reference genome. This *E. coli* strain differs from the Phase 1 Ancestor, REL1206, in seven positions (*topA, spoT K662I, glmU/atpC,pykF, yeiB, fimA* and the *rbs* operon) that were excluded from our analysis (Barrick et al. 2009; Tenaillon et al. 2012). We performed a first-round of variant detection using breseq in polymorphism mode on all of evolved populations relative to *E. coli* strain B REL606.

Following this first-step of the analysis, we generated a new, mutated reference sequence to represent each Phase 2 Founder using the gdtools APPLY command in breseq using the sequencing data available in Tenaillon et al. (2012). We then ran the breseq analysis again with respect to the Phase 2 Founder using the mutated references to verify mutation predictions, as described in Deatherage and Barrick (2014). Using gdtools available through breseq, we compiled the mutation information into readable tables and as an alignment file in PHYLIP format. A phylogeny was constructed using IQ-tree and the PHYLIP alignment as input (Nguyen et al. 2015).

### Statistical Analyses

All statistical analyses were performed in R v 4.0.2 (R Core Team 2019). For relative fitness results and statistical analysis, we first assessed normality of the data using the Shapiro-Wilk test and the variance with Levene’s test available through R. To statistically test for associations between the mutation patterns observed in Phase 2 and their initial adaptive pathway, we first built a distance matrix from the presence and absence matrix of Phase 2 mutations in R. Using the vegan package v 2.5-7 in R, we directly tested for associations between the distance matrix of Phase 2 mutations and the adaptive pathway or mutated codon with ANOSIM (Oksanen et al. 2020). We also built a Neighbor-Joining (NJ) tree based on the presence-absence matrix of accessory genes. To do so, we first calculated the Euclidean distances from the presence-absence matrix of the accessory genes using the *dist* function in R. We then built the NJ tree from the Euclidean distances using the ape package in R (Paradis & Schliep 2019). We calculated Dice’s similarity coefficient (DSC) for each pair of Phase 2 evolved populations using the *vegan* package. Dice’s coefficient of similarity was calculated using the same distance matrix used to build the NJ tree described above, as well as on a distance matrix containing only the fixed mutations. To identify genes or intergenic regions that were more frequently mutated in populations descended from one adaptive history but not the other, we built a 2 × 2 Fisher’s Exact Test for each mutated gene or intergenic region in R, for a total of 163 contingency tests.

## DATA AVAILABILITY STATEMENT

All high-throughput sequence data generated in this study have been submitted to the NCBI database and can be accessed through BioProject #######.

## ACKNOWLEDGEMENTS

We thank R. Gaut and the UC Irvine Genomics High-Throughput Facility for contributing to data generation. We also thank A. Martiny for helpful input on methodology. T. Batarseh was supported by the National Science Foundation Graduate Research Fellowship Program and the University of California Irvine President’s Dissertation Year Fellowship.

## Notes

### Competing Interest Statement

The authors have declared no competing interest.

