## Supplemental Figures for "Phenotypic and genotypic adaptation of *E. coli* to thermal stress is contingent on genetic background"

### Supplementary Figures

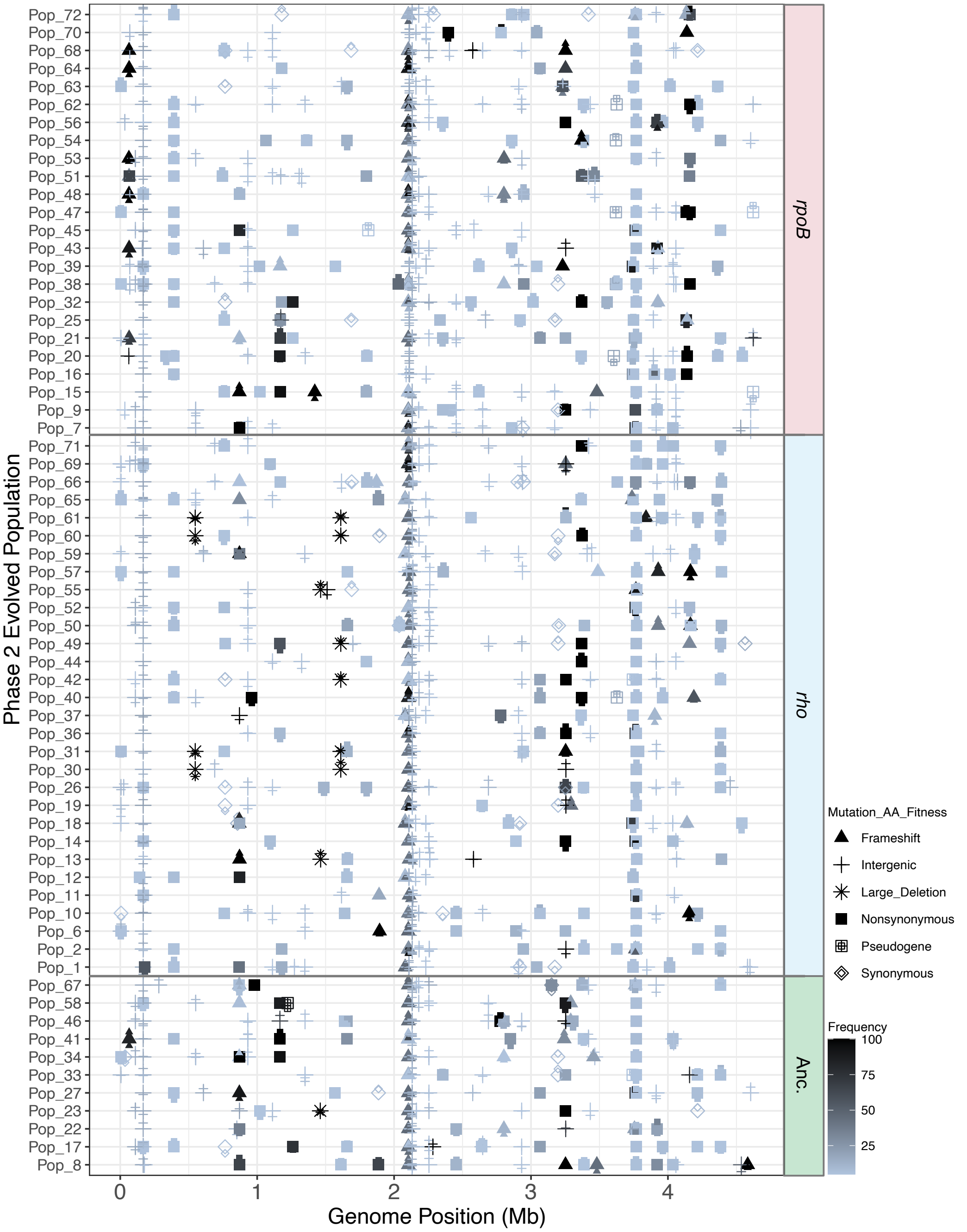


**Supplemental Figure 1:** All detected mutations at a frequency of 5% or higher in the Phase 2 evolved populations represented along the *E. coli* chromosome. Mutational types are distinguished by the shape of the point and the intensity of the color corresponds to the frequency that mutation was found in the population. Populations are grouped to the right according to their Phase 2 Founder historical background.

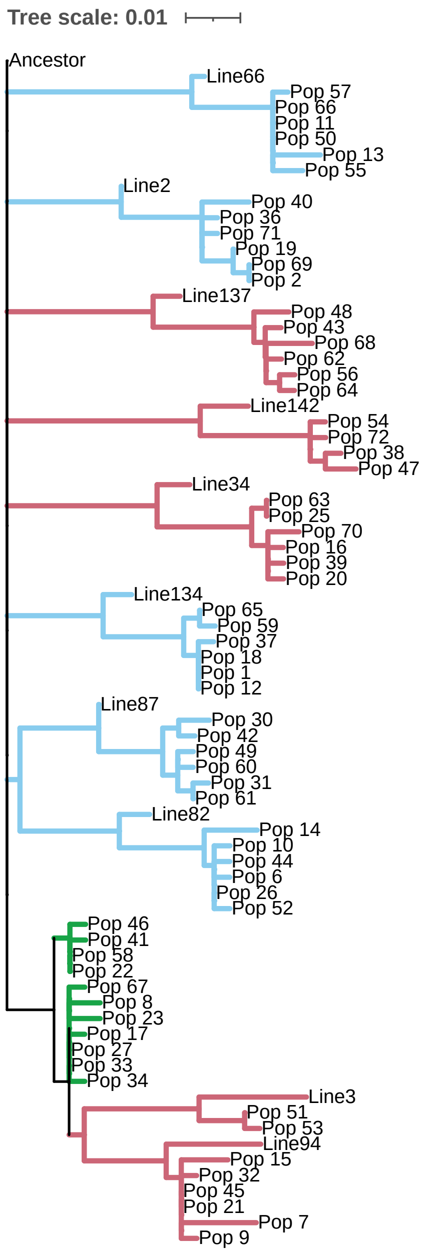


**Supplemental Figure 2:** Maximum likelihood phylogeny built from all mutation data across both phases of the experiment. Sequencing data from the ancestral *E. coli* REL606, the Phase 1 Founders, and the Phase 2 evolved populations were used to build the phylogeny. Branches are colored according to the Phase 2 Founder historical background: *rpoB* = red, *rho* = blue, and ancestral (REL1206, Phase 1 Ancestor) = green.

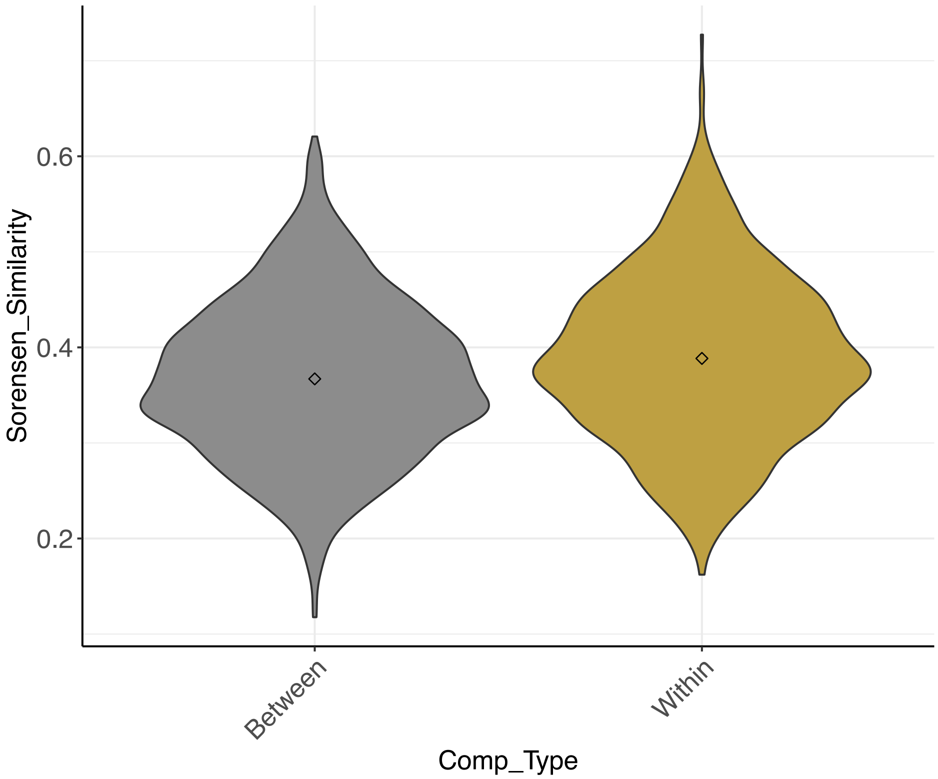


**Supplemental Figure 3:** Dice’s similarity coefficients calculated from total (5% or higher frequency) mutation data, separated by the type of pairwise comparison
